# Pleasant touch mediated by A*β*-afferents remains primarily dependent on tactile velocity

**DOI:** 10.64898/2025.11.29.690443

**Authors:** Thanh-loan Sarah Le, Dimitri Vinet, Gilles Bailly, Malika Auvray, David Gueorguiev

**Affiliations:** Sorbonne Université, CNRS, Institut des Systèmes Intelligents et de Robotique, F-75005 Paris, France; UCLouvain, Institute of Neurosciences, B-1200 Woluwe-Saint-Lambert, Belgium

**Keywords:** A*β*-afferents, CT afferents, pleasant touch, vibrotactile stroke, brush stroke, contrast effect, incongruence, virtual reality

## Abstract

Pleasantness of touch is classically attributed to C-tactile (CT) afferents, based on multiple evidence showing identical inverted-U curves as a function of velocities for pleasantness and CT activity. Recent research suggests that pleasant touch is not solely mediated by CT activation, pointing to a potential contribution of A*β*-afferents and cognitive processes, potentially through the mechanism of velocity inference. To investigate the contributions of A*β*-afferents, our study compared the illusion of vibrotactile apparent motion, which recruits almost exclusively A*β*-afferents, and brush stroking, which additionally recruits CT afferents. A first experiment investigated the impact of stroke contrast on the pleasantness of 3 cm/s strokes by mingling them with pleasant (1 and 10 cm/s) or less pleasant (0.5 and 30 cm/s) ones. A second experiment in virtual reality probed velocity inference from visual cue by testing how congruence between visual and tactile velocity affects perceived pleasantness. The results showed that the pleasantness of strokes targeting A*β*-afferents also follows an inverted U-curve and that 3 cm/s reference strokes remained unaffected by the speed of mixed strokes regardless of the condition. Even though differences in incongruence were not clearly perceived, discrepant visual velocities influenced the pleasantness of vibrotactile 1 cm/s strokes, for which the pleasantness ratings followed the visual rather than the tactile velocity. Overall, vibrotactile pleasantness can be mediated by A*β*-afferents, but most probably through velocity inference since only vibrotactile strokes were prone to visual illusion. Because not clearly detected, visual inferences could have played a stronger role through larger speed discrepancies.

## Introduction

Daily affective interactions strongly impact human emotional well-being and strengthen social bonds (Field, 2010; Hauser et al., 2019; Hertenstein et al., 2006). Gentle strokes on hairy skin are known to activate C-tactile (CT) afferents, following an inverted U-shaped sensitivity curve with greater activation between 1 and 10 cm/s that peaks at 3 cm/s (Ackerley et al., 2014; Löken et al., 2009; Olausson et al., 2010). Numerous studies showed this firing pattern to mirror the subjective ratings of pleasantness (Ackerley et al., 2014; Crucianelli et al., 2022; Löken et al., 2009, 2011; Luong et al., 2017; Triscoli et al., 2013). In line with the scientific results, humans naturally use velocities that strongly activate CTs when stroking their partner or child (Croy et al., 2016). Moreover, natural skin temperature (32°) elicits stronger CT responses than when the skin is cooled (18°) or heated (42°) (Ackerley et al., 2014). These multiple links between CT activity, pleasantness, and social touch have sparked the "Social Touch Hypothesis", which states that CT afferents constitute the neural basis of the emotional pathway of touch (Morrison et al., 2010; Vallbo et al., 1999).

Fast-conducting myelinated A*β*-afferents, which primarily convey discrimination abilities, are also activated during affective touch (Ackerley & Kavounoudias, 2015; Bolanowski et al., 1988; McGlone et al., 2014). Given that they do not show a straightforward correlation with the velocity of stroke, they have been thought to play at most a minor role in mediating social touch (Löken et al., 2009; McGlone et al., 2014). Nevertheless, recent articles have suggested a potential role of A*β*-afferents in conveying tactile pleasantness. A reduction of A-fiber function through an ischaemic nerve block has been shown to almost entirely abolish the perception of pleasantness (Case et al., 2023). Furthermore, ablation of the I-spinothalamic pathway, to which CT afferents project, eliminates the perception of pain and temperature, but not the pleasant sensations associated with the speed of stroking (Marshall et al., 2019).

The potential pathway, provided by A-fibers, could explain why apparent vibrotactile motion on the forearm can be experienced as pleasant (Huisman et al., 2016; Israr & Abnousi, 2011), although CT activity does not correlate with vibrotactile frequency and produces only a few impulses during low-frequency sinusoidal vibration (Ackerley, 2022; Bessou et al., 1971). The literature provide contrasted results on the role of Pacinian corpuscles, i.e. A*β*-afferents with peak sensitivity around 200 Hz, on pleasantness. While some studies suggest that they do play an important role in conveying pleasantness (Huisman et al., 2016), another one reported no effect of stroking velocity on pleasantness rating of vibrotactile apparent motion (Israr & Abnousi, 2011), which might come from the narrower range of velocities tested (5.7, 7, 9, 13, and 23 cm/s) in comparison with the typical span (0.3–30 cm/s) (Taneja et al., 2021). A recent study compared the pleasantness ratings elicited by continuous motion of a soft silicon end-effector on the skin and by apparent motion induced by successive taps of the silicon end-effector (Pehkonen et al., 2025). The obtained results showed that the apparent motion, which targeted CTs in a manner that did not provide a direct velocity cue, still induced an inverted U-shaped curve, suggesting that pleasantness stems from higher-order perception of velocity rather than from its direct encoding by CT afferents.

The pleasantness curve can still be observed even in the absence of touch; it can be triggered simply by observing someone else experiencing pleasant touch (Lee et al., 2018; Morrison et al., 2011; Walker et al., 2017; Willemse et al., 2016). This phenomenon shows the cognitive role played by the characteristics of a visually perceived stroke, which is in line with broader research on the cognitive inference of tactile velocity. For example, the cutaneous rabbit illusion, a tactile spatio-temporal illusion of motion, was shown to be shaped by prior expectations of speed (Goldreich, 2007). In addition, the type of texture can strongly bias the perceived tactile speed during passive touch (Delhaye et al., 2019). Thus, in the two aforementioned studies that used vibrotactile apparent motion, it is possible that participants relied on inferences about stroke speed when assessing pleasantness. The particularly difficult discrimination of tactile speed for vibrotactile apparent motion with a Weber fraction above 0.5 (Kohli et al., 2006; Lacôte et al., 2022) might explain why a wide range of velocities is needed to elicit pleasantness ratings that follow the classic U-shaped curve. Alternatively, pleasantness ratings could have been impacted by the reference points that the clearly unpleasant stimuli (0.3 and 30 cm/s) provide, a phenomenon called contrast effect. The pleasantness of 3 cm/s and 30 cm/s strokes is similar when each group is subjected to only one speed. This is in contrast to studies in which participants experience multiple speeds (Triscoli et al., 2014). Moreover, the order of stroke delivery between the palm and forearm was found to affect the curve of pleasantness elicited during evaluations of the forearm’s affective response (Löken et al., 2011).

The pleasantness of affective touch is also shaped by cross-modal interactions stemming from the multisensory context of interactions (Seinfeld et al., 2022; Spence, 2022; Sun et al., 2024). Spatio-temporally congruent visual and tactile feedback in virtual reality has been shown to increase the feelings of pleasantness compared to single visual stimulation, whether it involves a real interpersonal touch (Sun et al., 2024) or ultrasonic mid-air haptic stimulation (Seinfeld et al., 2022). The speed of visual distractors has also been shown to influence the perceived speed of moving tactile gratings (Bensmaïa et al., 2006), which might affect ratings of tactile pleasantness in multisensory contexts. Indeed, the pleasantness of congruent visuo-tactile strokes in virtual reality was shown to follow an inverted U-shaped curve, while shifting the velocity of the visual stimulus compared to the tactile stimulus modifies the curve (Haraguchi & Kitazaki, 2022).

The effect of stroke velocity on perceived touch pleasantness and its correlation with CT-afferents activity is a well-established phenomenon. Nevertheless, recent evidence suggests a potential role of both A-fibers activity and cognitive inference, which relies on either priors or discrepant multisensory cues. In this context, our study aims to assess the perceived pleasantness of dynamic strokes that target almost exclusively A*β* afferents by using the illusion of vibrotactile apparent motion, and to compare it with the one elicited by brush strokes triggering both A*β* and CT afferents. In addition, our study investigates the influence of two cognitive mechanisms: the contrast effect and visual velocity inference.

In a first experiment, the perceived pleasantness of a reference stroke at 3 cm/s was compared to slower (0.5 and 1 cm/s) and faster (10 and 30 cm/s) velocities. The 3 cm/s stroke was used as the reference point, as it represents the highest level of pleasantness and the strongest CT activity. The stroke was either delivered by apparent vibrotactile motion or by a soft brush moved by a robotic arm. We hypothesized that stroke pleasantness at 3 cm/s would be enhanced by concomitant notoriously unpleasant velocities (0.5 cm/s or 30 cm/s), compared to concomitant moderate strokes (1 cm/s or 10 cm/s) that are usually considered rather pleasant.

A second experiment with the same apparatus was conducted in virtual reality to investigate conditions in which visual and tactile brush strokes / vibrotactile strokes were congruent or incongruent. The velocity of the tactile strokes was either 1 cm/s or 10 cm/s, while several visual velocities were delivered, defined as multiplicative factors of the tactile velocities (0.5, 0.75, 1, 1.25, and 1.5). A perception test was added to probe whether participants noticed the visuo-haptic incongruence in the stimuli. We expected the visual cues congruent with touch to enhance its perceived pleasantness, and that vibrotactile strokes, due to their illusory nature, would be more impacted by visual discrepancy. Across all experiments, brush strokes were predicted to yield higher pleasantness ratings, reflecting additional CT afferent activation.

## Materials and Methods

### Participants

24 participants completed the first experiment (13 women, 11 men, mean age 25.7 yr, *SD* = 2.2). They were stimulated both by a brush and by a vibrotactile device delivering apparent haptic motion at various velocities. In the second experiment, two groups were formed to avoid excessively long experiment durations in virtual reality. A "Brush" group of 24 participants (11 women, 13 men, mean age 25.6 yr, *SD* = 3.5) was stimulated with the brush and a "Vibrotactile" group of 24 participants (14 women, 10 men, mean age 28.8 yr, *SD* = 7) experienced vibrotactile apparent motion. The experiment was approved by the Research Ethics Committee at Sorbonne Université, under the approval CER-2023-LE-VS and was conducted in accordance with the guidelines of the Declaration of Helsinki for research involving human participants.

### Experiment 1: Contrasting velocities of Vibrotactile and Brush strokes

#### Apparatus and Stimuli

##### Brush stimulation

A robotic arm (Franka Emika Research 3, Franka Robotics, Inc.) was programmed to perform physical strokes (Figure 1). The stroke velocities were set at 0.5, 1, 3, 10, and 30 cm/s. A 40 mm soft brush (synthetic brush, Blue Line Leonard), known to efficiently stimulate CT afferents (Ackerley et al., 2014; Löken et al., 2009), was attached to the end-effector of the robot. The force applied to the participant’s forearm was calibrated to 0.4 N (Löken et al., 2009) using a force sensor (Mini45-E, ATI Industrial Automation) connected to the robot’s end effector. In practice, the force exerted by the soft brush was displayed in real time on a computer screen and the dynamic strokes on the participant’s forearm (5 cm/s) were fine-tuned to apply a normal load of approximately 0.4 N during the stroke. During the stimulation, the brush, which was inclined at 45°, reached the skin following a Bezier curve, stroked the latero-ventral side of the participant’s forearm across 9 cm (proximal-to-distal), before rising again following again a Bezier curve (see Figure 1). The stroke ended 3 cm before the wrist.

**Figure 1.**
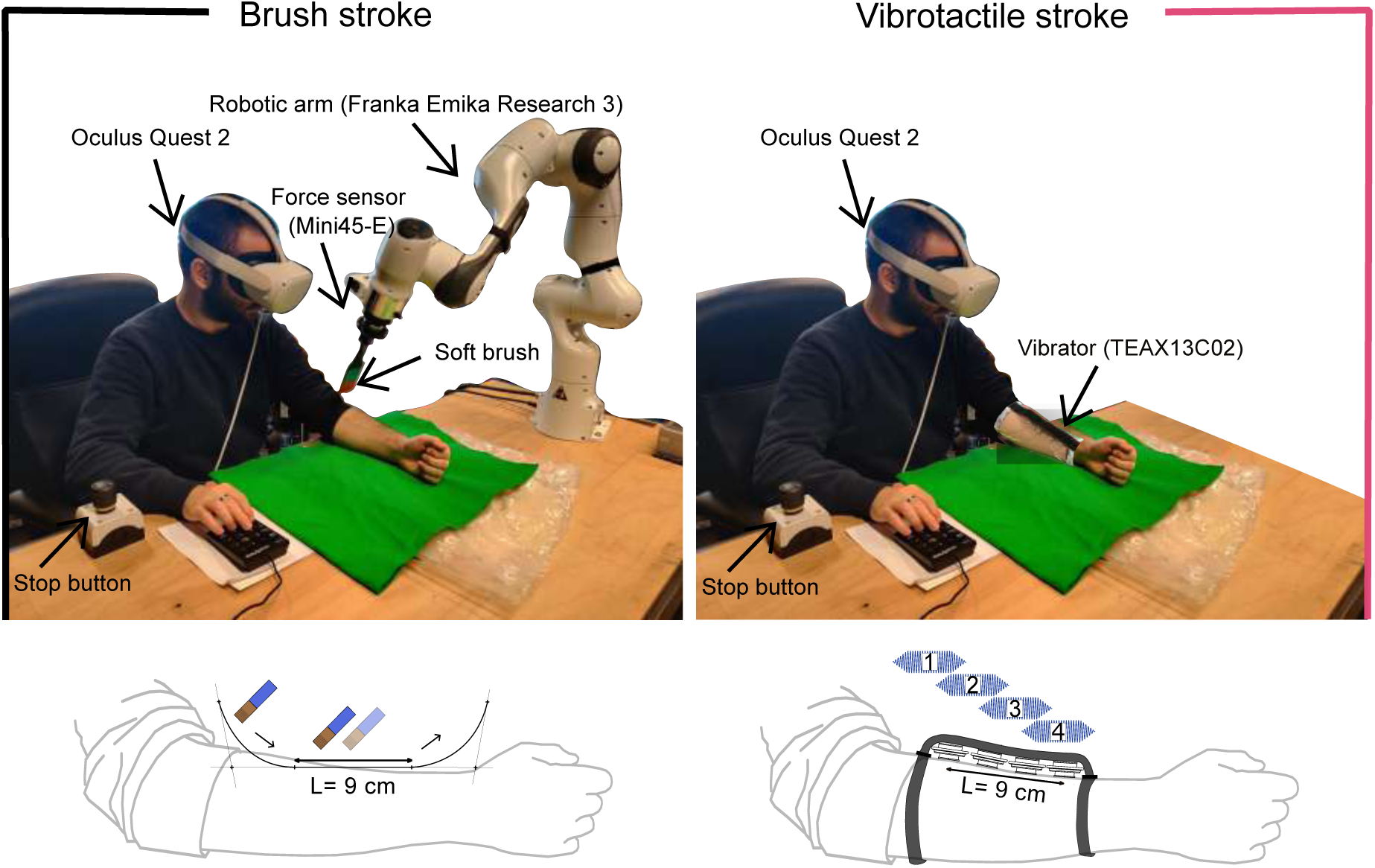
Experimental setup. Brush strokes were delivered using a brush-mounted robotic arm and Vibrotactile strokes by four sequentially activated vibrators producing apparent motion. In Experiment 1, participants received both stimulations in alternating block sessions. In Experiment 2, the "Brush" group received visuo-tactile brush strokes, while the "Vibrotactile" group received visual brush strokes in synchrony with vibrotactile strokes. The visual feedback was provided by an Oculus Quest 2 headset. An emergency stop button was placed between the participant and the experimenter. See S1 to see a video showing congruent visuo-tactile brush strokes in the first person perspective (Experiment 2). The user appearing on the photo belongs to study’s authorship.

##### Vibrotactile apparent motion

Four voice-coil vibrotactile actuators (TEAX13C02, Tectonic Elements, UK), identical to (Israr & Abnousi, 2011), generated vibrotactile apparent motion to create the vibrotactile strokes. The vibrotactile stroke velocities were set at 0.5, 1, 3, 10, and 30 cm/s. We chose to use four voice coils since this number is considered optimal to provide a convincing illusion (Nunez et al., 2020). The vibrators were placed lengthwise 3 cm apart (center-to-center distance) with a plaster on the lateral side of the participant’s forearm that was not stroked by the brush. (Figure 1). The most distal actuator was placed 3 cm above the wrist. Sinusoidal 120 Hz vibrations were recorded on wav. files and processed by a microcontroller STM32G474 with 4kHz sampling frequency, a 9 bit PWM resolution. Vibrotactile strokes resulted from the sequential activation of the four actuators (proximal-to-distal) (Figure 2A). The vibrotactile apparent motion relies on three parameters (in seconds): Duration of Stimulation (*DoS*); Stimulus Onset Asynchrony (*SOA*), i.e., the time delay between the actuator’s activation, and Total Stimulation Time (*TST*) (Israr & Poupyrev, 2011a; Kirman, 1974). The tactile velocity is computed as the ratio between the length of the stimulation (from first to last vibrator) and *TST* (see Appendix). The 120 Hz vibration was used, as in a previous study on apparent haptic motion (Sherrick & Rogers, 2010). Each vibration was ramped at onset and offset to reach the maximum intensity at 20% of its dura-tion, to smooth transitions. The vibrotactile signal was verified with an accelerometer (PCB Piezotronics Accelerometer 352A24) glued to the forearm’s skin 1 cm away from one actuator. (Figures 2B and 2C).

**Figure 2.**
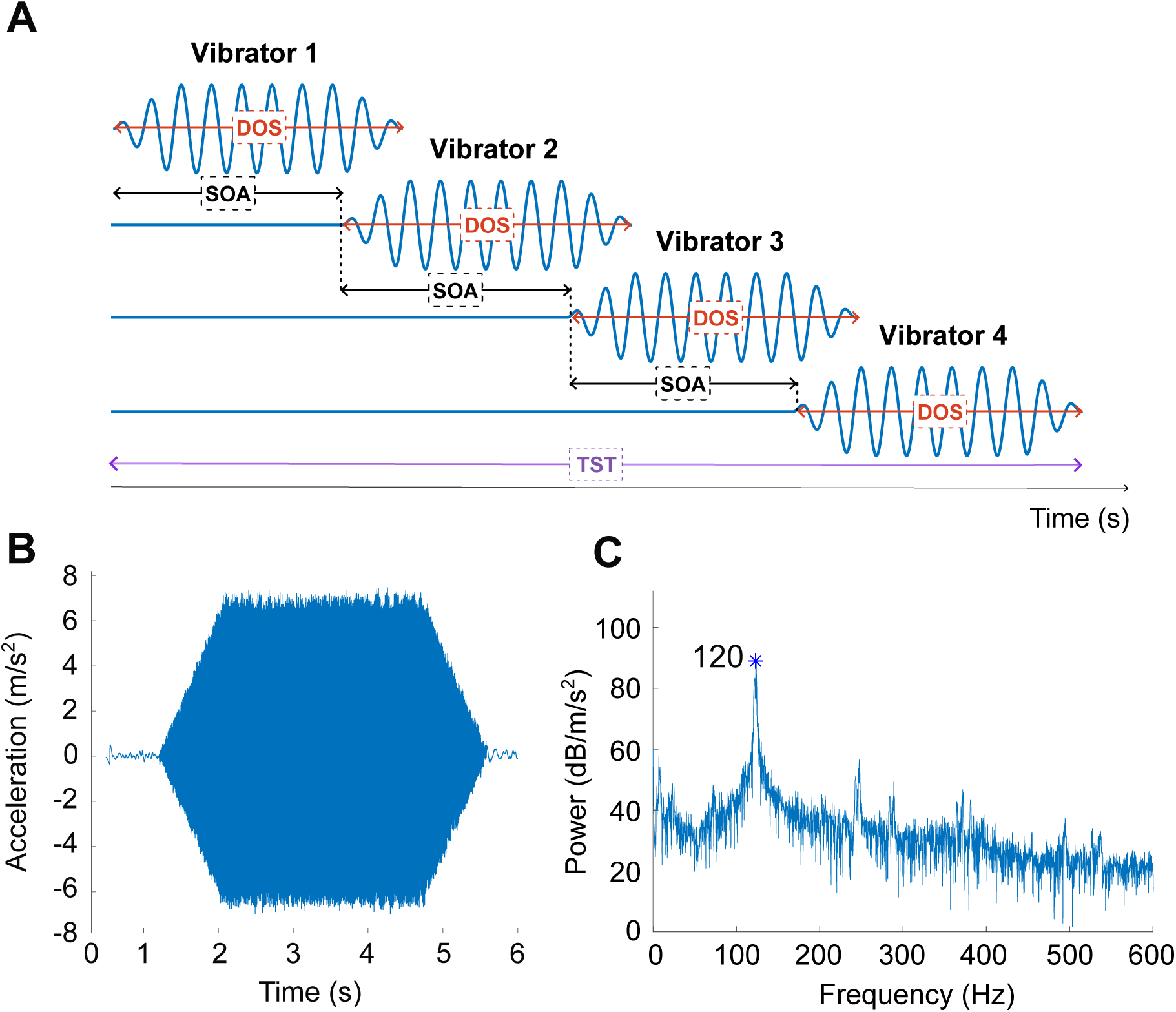
**A)** Vibrotactile apparent motion generated by four actuators vibrating at 120 Hz and defined by its Duration of Stimulation (*DoS*), Stimulus Onset Asynchrony (*SOA*) and Total Stimulation Time (*TST*). **B)** Measurement from an accelerometer located 1 cm away from a single vibrator placed 1 cm from it on the forearm skin **C)** Fast Fourier Transform showing the accuracy of the 120 Hz vibration.

#### Experimental procedure

All the participants were given the same standardized instructions and they were informed that all stimuli were innocuous. They were seated comfortably, provided with noise-cancelling headphones and the two stimulation sites were covered by a curtain. The participants’ forearms were comfortably positioned on a vacuum pillow filled with rice; one arm wore the vibro-tactile setup, while the other was stimulated by the brush (counterbalanced across participants).

##### Calibration phase

The participants started the study with a calibration phase that consisted in: (1) positioning the brush on the forearm’s skin so that the stroke can be performed with identical starting point and stimulation distance as the vibrotactile stroke; (2) calibrating the robot to apply a 0.4 N normal force during stroking at 5 cm/s, a velocity that is not used in the main experiment to avoid any influence; (3) equalizing the intensity of the vibrotactile stroke (5 cm/s) to the intensity experienced during the brush stroke. In (3), participants adjusted the intensity of vibrotactile strokes by using a Visual Analogue Scale (VAS) displayed on the computer in front of them (mild 0 - 100 intense, step = 5), until they perceived it as equivalent to the brush stimulation (Jones & Tan, 2013). Six repetitions, in which the initial amplitude was randomly set, were performed. The point of subjective equality for each participant was computed as the mean intensity and was subsequently used in the main experiment.

##### Pleasantness and intensity ratings

After the calibration phase, participants completed alternating blocks between brush and vibrotactile strokes. Two curtains prevented them from seeing the strokes, and they were instructed to keep their arms and hands still. At the end of each trial, they were instructed to rate the pleasantness and intensity evoked by the stroke using a VAS (from 0 to 100, step = 5) displayed on a laptop monitor, with the endpoints marked as “unpleasant” to “pleasant” for the pleasantness rating and “mild” to “very intense” for the intensity rating. One interval tick was placed at the middle of the VAS to prevent a potential bias to the left or right, but no other value was displayed. The starting position of the cursor was random in each trial. Participants used a joy-con (Switch, Nintendo Inc.) to move the slider and validate their answers. The Joy-con was operated by the arm equipped with the vibrotactile device. After each trial, a 15-second break took place for fatigue recovery of CT afferents (Vallbo et al., 1999) and potential vibrotactile adaptation (Erp, 2002).

The experiment consisted of eight blocks corresponding to combinations of the two stimulation type (brush and vibrotactile) and the four possible companion velocities. The presentation order was counterbalanced across participants. In each block, one companion velocity (0.5, 1, 10, or 30 cm/s) and the reference velocity (3 cm/s) were pseudo-randomly used six times each, resulting in a total of 96 strokes per participant. The reference stroke was thus presented more frequently than the other velocities. A single 5-minute break was given halfway through the experiment, which lasted approximately 80 minutes.

### Experiment 2: Visuo-haptic perception of stroke pleasantness

#### Apparatus and Stimuli

The experimental setup and the intensity of vibrotactile strokes were identical to Experiment 1, and the applied force of 0.4 N was calibrated with the same procedure.

##### Virtual reality scene

The virtual scene was programmed using Unity 3D and rendered with an Oculus Quest 2 (see S1). The virtual environment consisted of a room with a table, chairs, a sofa, and a lamp. During the experiment, the participants could see a virtual arm on the table from a first-person perspective in the same position as their physical arm. The avatar had a neutral skin colour (gray), was neutrally dressed in jeans and a white t-shirt, and displayed no distinctive features related to either male or female gender. The virtual stroke was performed by an animated brush. The incongruence between the visual and the tactile strokes was achieved by modifying the speed of the visual stroke. The stroking distance was also modified to keep the same duration of stimulation and thereby avoid an obvious mismatch between the start or end of the tactile stroke and its visual counterpart. The changes in distance were kept between 0 and 4.5 cm (see Figure 3), below the visuo-tactile two-point discrimination threshold (Caballero & Rombokas, 2019). To prevent participants from learning the cues, the starting point was randomized with the constraint that the virtual brush would stay above the wrist.

**Figure 3.**
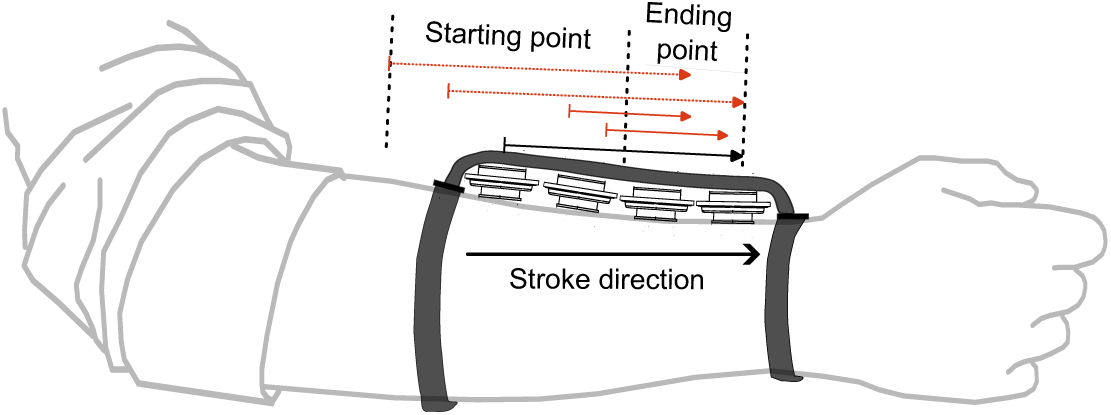
Illustration of the distances covered by the visual (red arrows) and tactile (black arrow) strokes in Experiment 2. The starting point for the visual strokes was positioned to ensure that the offset between the beginning of the visual and tactile strokes would never exceed 4.5 cm.

##### Stimuli

Participants were simultaneously presented with a tactile and visual stroke on the forearm. The tactile stroke had a speed of either 1 cm/s or 10 cm/s. For tactile strokes at 1 cm/s, the velocities of visual strokes could be 0.5, 0.75, 1, 1.25, and 1.50 cm/s; for tactile strokes at 10 cm/s, the velocities of visual strokes were 5, 7.5, 10, 12.5, and 15 cm/s. Thus, four visuo-haptic combinations were incongruent for each tactile speed. The possible distances of visual strokes were 4.5, 6.75, 9, 11.25, 13.5 cm, and the constant distance of the tactile strokes was 9 cm.

#### Experimental procedure

##### Pleasantness and intensity ratings in virtual reality

At the start of the experiment, the participants were asked to spend 30 seconds playing with their virtual arm and fingers to familiarise themselves with them. They were instructed to keep their arms and hands still while observing the stimulated forearm during each trial. The procedure for providing ratings was the same as in Experiment 1 (see Pleasantness and intensity ratings), except that participants used a numeric pad to answer because the Joy con was not compatible with the virtual reality setup. The first task included 10 different types of strokes (2 *Tactile velocities* x 5 *Visual velocities*) presented in random order and repeated 3 times (30 trials in total). The task took approximately 25 minutes to complete.

##### Congruence perception test

After a 10-min break, participants completed the congruence perception test. In this second task, they indicated whether the speed of visual and tactile strokes was congruent (Yes/No task), using the dialogue box shown in the virtual scene. The same strokes as in task one were played. Congruent strokes were repeated six times (12 strokes) and incongruent strokes three times (24 strokes). The task lasted approximately 25 minutes.

### Statistical analyses

In all experiments, averaged scores across repetitions were analysed with Linear Mixed Models in RStudio (4.1.3). Linear and quadratic effects were tested, and model selection was based on Likelihood Ratio Tests. The Linear Mixed Models were fitted using maximum likelihood estimation with a Gaussian error distribution. Shapiro–Wilk tests indicated that the raw data in each analysis were not normally distributed; however, the distribution of model residuals was normal. In Experiment 1, the dependent variables were the magnitude of perceived *Pleasantness* and *Intensity*, and two categorical independent variables were *Stimulation* and *Companion velocity*. In task 1 of Experiment 2, dependent variables were the same and independent variables were *Stimulation*, *Tactile velocity*, and *Visual velocity*. In task 2, the dependent variable was the *Proportion of congruent answers*. Type III Wald Chi-square tests were used to test independent variables in Experiments 1 and 2. Post-hoc pairwise comparisons were conducted on significant fixed effects between estimated marginal means by using the package “emmeans” of RStudio. P-values were adjusted for multiple comparisons using either Tukey’s method for complete pairwise comparisons and Holm’s method for selected contrasts. To test whether fixed-effect slopes (estimates *β*) differed significantly from zero, t-statistics were computed from the estimated coefficients using Satterthwaite’s method ("lmerTest" package). This method has the advantage of adjusting the degree of freedom according to random factors. Finally, Spearman’s correlation tests were performed between pleasantness and intensity ratings.

## Results

### Experiment 1

#### Stable perceptual pleasantness of 3 cm/s tactile stroking

The Wald analysis conducted on the Linear Mixed Model (LMM) (*Stimulation* and *Companion velocity*) showed a significant main effect of *Stimulation* (*χ*^2^(1) = 23.53, *p <* 0.0001) on *Pleasantness* of 3 cm/s strokes, while there was no significant effect of *Companion velocity* (*χ*^2^(3) = 4.23, *p* = 0.24). Moreover, no significant interaction effect between *Stimulation* and *Companion velocity* was found (*χ*^2^(3) = 3.76, *p* = 0.29). Post-hoc test on *Stimulation* revealed that brush strokes (*M* = 67.4, *SD* = 16.7) were significantly more pleasant than vibrotactile strokes (*M* = 60.1, *SD* = 16.7) (Tukey-adjusted pairwise comparisons: *n* = 96, *p <* 0.0001) (Figure 4A). Finally, we controlled for the impact of the blocks’ presentation order since each block features a distinct companion velocity, and no significant effect was observed (*χ*^2^(7) = 5.60, *p* = 0.58).

**Figure 4.**
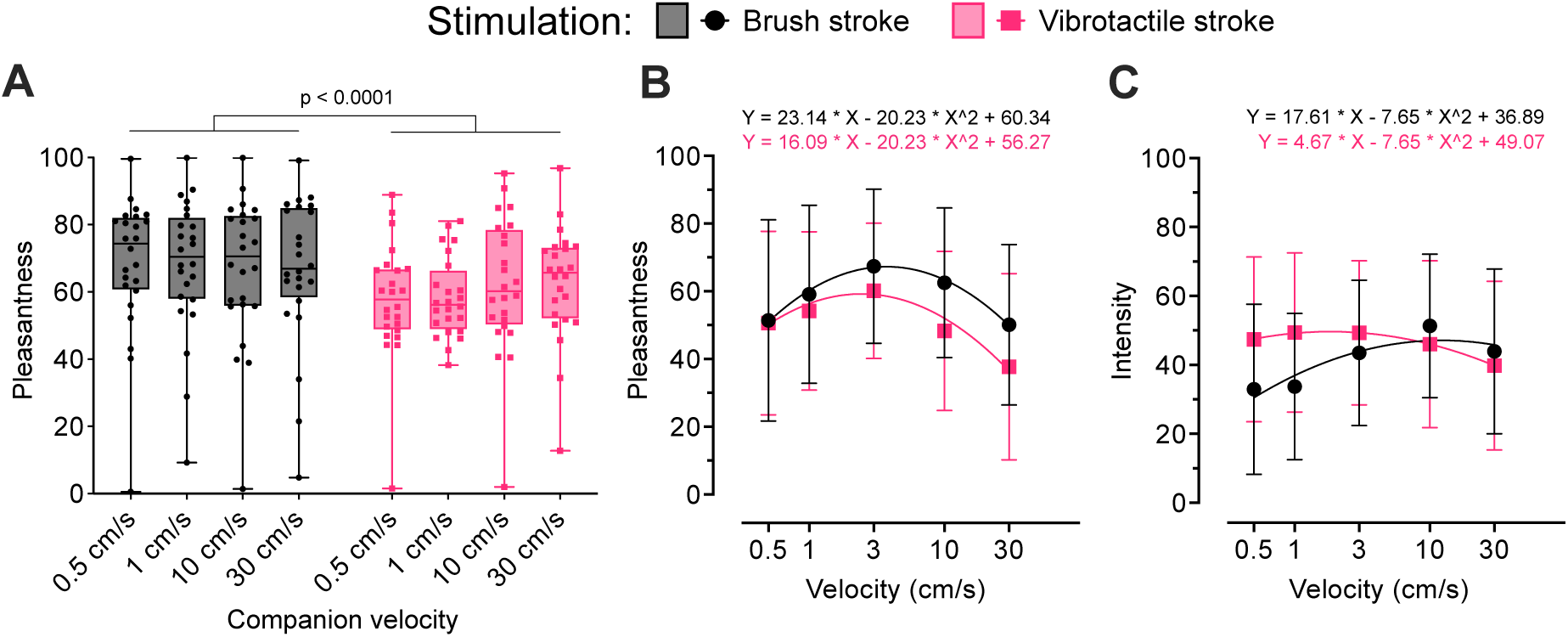
**A)** *Pleasantness* rating (median, interquartile range and min to max) of strokes at 3 cm/s averaged across participants’ mean ratings as a function of *Companion velocity* and *Stimulation*. **B)** *Pleasantness* of strokes as a function of *Velocity* and *Stimulation*. Lines show predictions from the full linear mixed model. **C)** Same analysis for *Intensity*.

#### Inverted U-shaped pattern of pleasantness and intensity

We analysed whether the pleasantness and intensity ratings exhibited an inverted U-shaped curve for the two types of stimuli. To that end, four LMMs were computed for pleasantness and intensity ratings of brush strokes and vibrotactile strokes as a function of velocity. The factor velocity was considered on a logarithmic scale as in previous studies (Ackerley et al., 2014; Löken et al., 2009. In each case, using models based on quadratic functions provided a better fit to the data than linear functions (Likelihood Ratio Test, *p−values <* 0.001).

With this model, we then investigated the difference in perceived *Pleasantness* as a function of the type of stimulation and velocity (Figure 4B). To that end, we built a global LMM (*Stimulation* and log(*Velocity*)) that incorporates quadratic effect terms. We compared a model including an interaction between curvature and stimulation type with a reduced model assuming a shared curvature. The Likelihood Ratio Test did not reveal a statistical difference between the two models, suggesting that differences in fitted coefficients do not provide evidence for different curvature profiles between stimulation types (*χ*^2^(1) = 0.19, *p* = 0.65). t-statistics computed on the estimated coefficients of the LMM showed that log(*Velocity*) has a significant effect on *Pleasantness* (*β* = 16.09, *SE* = 3.54, *p <* 0.001) and that the quadratic term is significant (*β* = *−*20.23, *SE* = 2.39, *p <* 0.001), confirming the non-linear relationship. Moreover, the main effect of *Stimulation* was marginal (*β* = 4.08, *SE* = 2.16, *p* = 0.06) but the interaction effect between log*(Velocity)* and *Stimulation* was significant (*β* = 7.05, *SE* = 2.99, *p* = 0.019) (Figure 4B). When the strokes were made by a brush, the level of pleasantness increased with velocity up to a maximum reached at 3.69 cm/s before decreasing for higher velocities. For vibrotactile strokes, the initial increase is less pronounced and the maximum is reached at the lower velocities of 2.44 cm/s (Figure 4B).

The same analysis was repeated on perceived *Intensity* using a global LMM that does not include an interaction between (*Stimulation* and log*(Velocity)*) (*χ*^2^(1) = 0.21, *p* = 0.64) (Figure 4C). t-statistics computed on the estimated coefficients of the LMM showed that the quadratic term log(*Velocity*) has a significant effect on *Intensity* (*β* = 4.66, *SE* = 3.18, *p* = 0.0004), as well as *Stimulation* (*β* = *−*12.07, *SE* = 1.95, *p <* 0.0001). Moreover, there was a significant interaction effect between log*(Velocity)* and *Stimulation* (*β* = 12.94, *SE* = 2.69, *p <* 0.0001) (Figure 4C). The quadratic model for Brush stroke stimulation shows a maximum peak at 14.2 cm/s, whereas that of Vibrotactile stroke reaches its peak at the much lower velocity of 2.02 cm/s (Figure 4C).

An additional analysis showed a weak, but still significant, association between the perceived pleasantness and intensity in the case of brush strokes (Spearman’s test: *r* = 0.26, *p <* 0.0001) (Figure 5). In the case of vibrotactile strokes, we observed a weak correlation (Spearman’s test: *r* = 0.14, *p* = 0.03), but this effect did not survive Bonferroni correction for multiple comparisons (*p_ad j_* = 0.06) (Figure 5).

**Figure 5.**
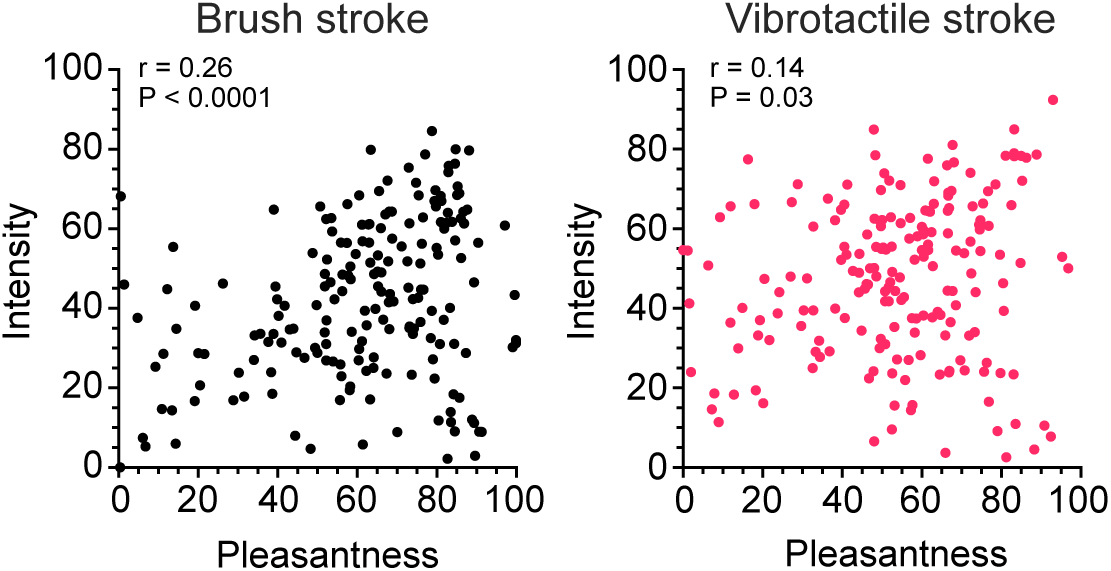
Correlation level between perceived *Pleasantness* and *Intensity* for brush and vibrotactile stimulations in Experiment 1. Data points represents the averaged participants’ values across velocities.

Overall, our results confirm that vibrotactile strokes exhibit a pleasantness rating that follows a classic inverted U-shaped curve. Ratings are similar to those elicited by classic brush strokes at low velocities but become significantly lower when velocity increases. Results also showed that the judgment of 3 cm/s strokes is not affected by the velocity of the other strokes present in a given experimental block, regardless of the type of stimulation. Brush strokes were rated more pleasant than vibrotactile ones, a difference that grew with velocity.

### Experiment 2

#### Visual inference on pleasantness

We analysed whether the pleasantness of tactile stroke varies as a function of visual velocity in both stimulation groups. For "Brush" group, the t-statistics computed on the estimated coefficients of the LMM (*Tactile velocity* and *Visual velocity*) showed no effect of *Visual Velocity* (*β* = 1.59, *SE* = 3.37, *p* = 0.64) nor *Tactile velocity* (*β* = 2.38, *SE* = 5.06, *p* = 0.64) on the perceived *Pleasantness* (Figure 6A). For the "Vibrotactile" group, the t-statistics showed a significant effect of *Visual Velocity* (*β* = 7.92, *SE* = 3.29, *p* = 0.017), but no effect of *Tactile velocity* (*β* = 7.01, *SE* = 4.93, *p* = 0.64) on the perceived *Pleasantness*. Moreover, the t-statistics showed a significant interaction between *Tactile velocity* and *Visual Velocity* (*β* = *−*8.16, *SE* = 3.31, *p* = 0.014) (Figure 6B). Indeed, a significant visual influence occurred only for slow vibrotactile strokes at 1 cm/s.

**Figure 6.**
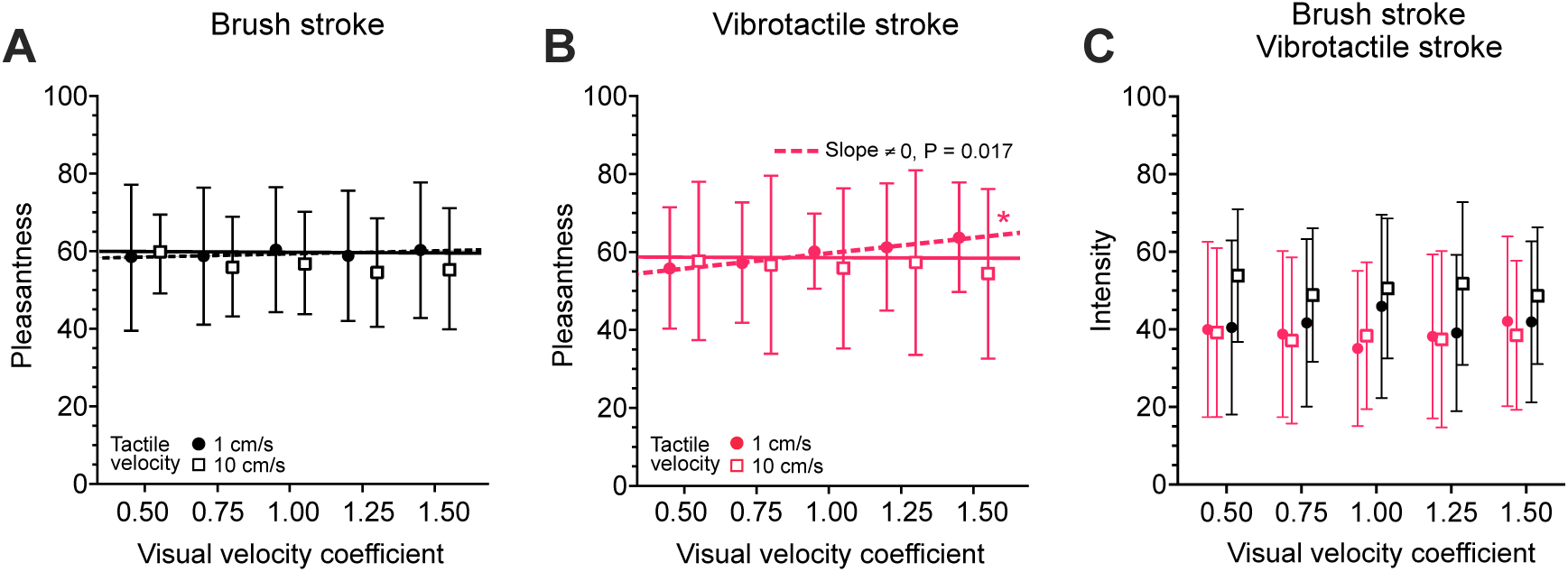
**A)** *Pleasantness* ratings (mean and SD) as a function of *Visual velocity* transformed by a coefficient for the two tested tactile velocities for the "Brush" group. Data points represent averaged value across participants at the corresponding visuo-tactile velocity. Lines show predictions from the full linear mixed model. An horizontal jitter was applied (x = 0.025). **B)** Same graph for the "Vibrotactile" group. **C)** *Intensity* ratings (mean and SD) as a function of *Visual velocity* transformed by a coefficient for the two tested tactile velocities. Both the "Brush" (black) and "Vibrotactile" (pink) groups are represented. A horizontal jitter was applied (x = 0.07).

Finally, the t-statistics computed on the estimated coefficients of the LMM (*Tactile velocity* and *Visual velocity*) on the perceived *Intensity* revealed no significant effect of both factors whatever the type of stimulation (Figure 6C).

To evaluate the strength of the visual impact (Figure 6B), we transformed the data from Experiment 1 to a linear scale instead of the logarithmic one (Figure 7), and we computed linear approximations of the pleasantness curve at 1 cm/s and 10 cm/s by using the tangent of the quadratic function. For vibrotactile strokes at 1 cm/s, the incongruent visual velocities created a slope of *β* = 7.92, similar to the slope elicited by tactile velocity in Experiment 1 (*β* = 6.44). For brush strokes at 1 cm/s, incongruent visual velocities could not elicit a significant change in pleasantness (*β* = 1.59 vs. *β* = 10.6 in Experiment 1). For both types of tactile strokes at 10 cm/s, the pleasantness was unaffected by incongruent visual velocity. However, the slopes computed from Experiment 1 also showed very little decrease in pleasantness (Brush stroke: *β* = *−*0.79 and Vibrotactile stroke: *β* = *−*1.02). Thus, a larger span of visual velocities might have been needed to elicit a visual for 10 cm/s.

**Figure 7.**
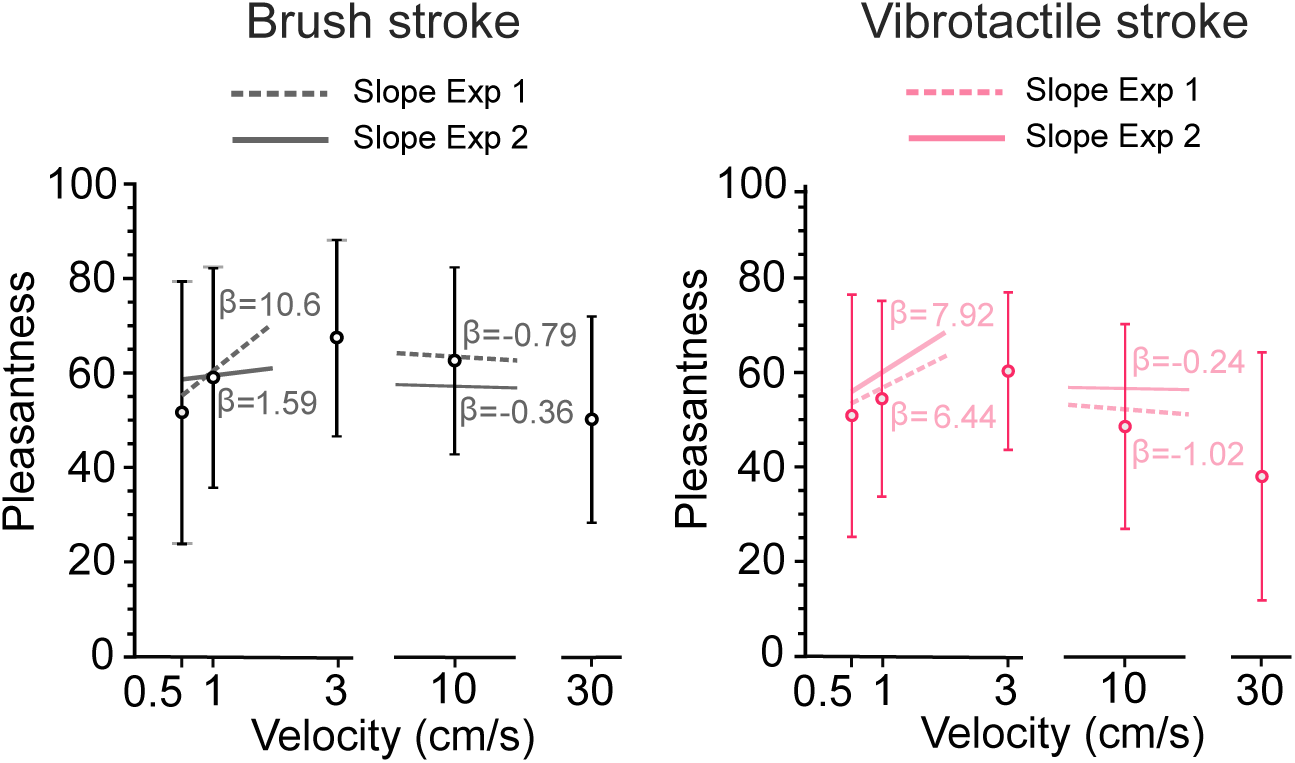
*Pleasantness* ratings (mean and SD) of brush and vibrotactile stroke stimulations in Experiment 1, shown on a linear velocity scale (cm/s). The x-axis is discontinuous to enable a clearer view of the data. Dashed lines correspond to the slope at 1 cm/s and 10 cm/s from the quadratic model fits in Experiment 1 and solid lines represent model fits in Experiment 2.

#### Detection of incongruence

After each trial, participants were asked to judge whether the visuo-tactile strokes they received were congruent or incongruent. For both types of stimulation, Wald tests showed a main effect of *Tactile velocity* (*p−values <* 0.0001) and *Visual velocity* (*p−values <* 0.01) on the *Proportion of congruent answers*, but failed to detect a significant interaction effect (*p − values >* 0.05). Post-hoc Tukey-adjusted pairwise comparisons on *Tactile velocity* (*n* = 432*, p − values <* 0.0001) showed that participants rated tactile 10 cm/s strokes as more congruent (Brush stroke: *M* = 0.79, *SD* = 0.07; Vibrotactile stroke: *M* = 0.84, *SD* = 0.09) than strokes moving at 1 cm/s (Brush stroke: *M* = 0.63*, SD* = 0.1; Vibrotactile stroke: *M* = 0.56, *SD* = 0.09). The only significant differences in perceived incongruence were found through Holm-adjusted pairwise comparisons between 0.5 cm/s and 1 cm/s for both stimulations types (*p −values <* 0.01) (Figures 8A and 8B) and between 1 cm/s and 1.25 cm/s for vibrotactile strokes (*p* = 0.042) (Figure 8B). Thus, in most cases, incongruent strokes were not perceived differently from congruent strokes.

**Figure 8.**
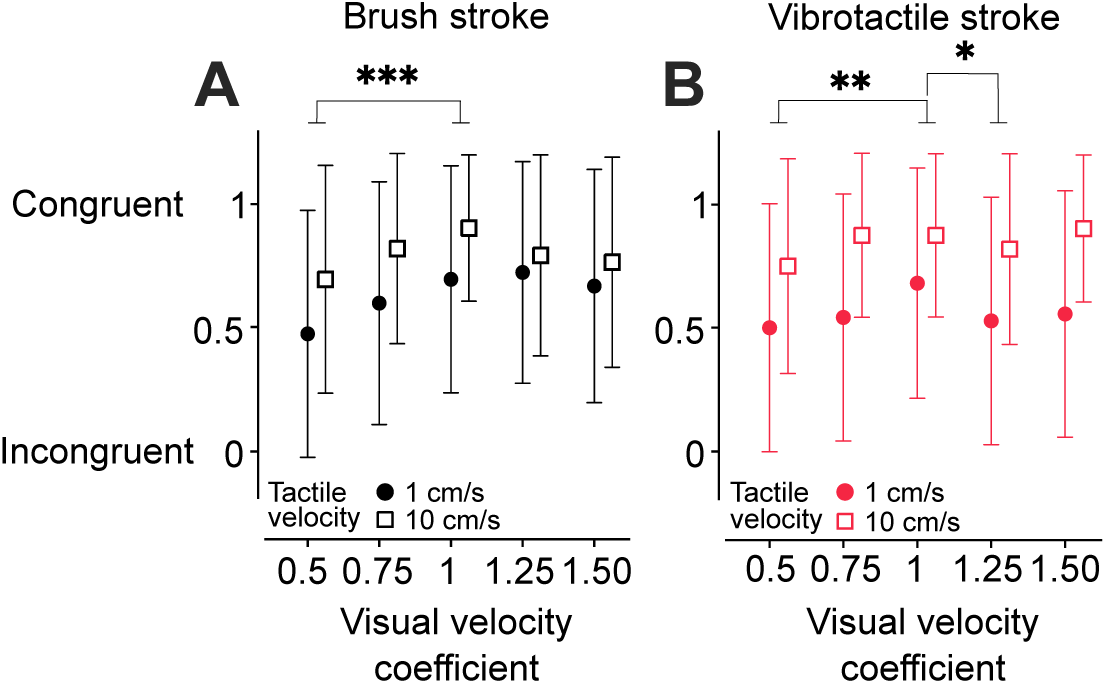
**A)** *Proportion of congruent answers* (mean and SD) as a function of *Visual velocity* transformed by a coefficient for the two tested tactile velocities for the "Brush" group. **B)** Same graph for the "Vibrotactile" group. Data points represent averaged values across participants at the corresponding visuo-tactile velocity. A horizontal jitter was applied (x = 0.025).

## Discussion

The present study aimed to investigate the respective contributions of afferent types and cognitive context in pleasantness of touch by comparing strokes that selectively activate A*β* through vibrotactile apparent motion and CT + A*β* afferents with a classic brush stroke. This approach was motivated by the potential role of A*β* afferents in affective touch, as well as cognitive processes that may combine with CT afferents-driven pleasantness. Our results confirmed that pleasantness of vibrotactile strokes also follows an inverted U-shaped curve as a function of velocity (Figure 4B), which supports the idea that A*β* afferents also mediate pleasantness of touch. No specific effect of the contrast induced by different companion velocities was observed for vibrotactile and brush strokes at 3 cm/s (Figure 4A), but discrepant visual velocities in virtual reality significantly influenced the pleasantness of vibrotactile strokes with constant 1 cm/s velocity (Figure 6B and Figure 7). This finding suggest that the pleasantness induced by A*β*-strokes is more prone to alteration through visual inference.

Vibrotactile strokes were rated as less pleasant than brush strokes, though not at the slowest strokes (0.5 cm/s and 1 cm/s), which elicited a pleasantness comparable to brush strokes. The consistent interaction between stimulation type and velocity observed across the study suggests different coding mechanisms between vibrotactile and brush stimulations. Overall, our study confirmed the pivotal role of velocity in perceived pleasantness even in the absence of CT afferents, but it also showed a specific resilience of CT afferents to discrepant visual strokes that highlights a dominant role of CT afferent pathways.

### Differences between brush strokes and vibrotactile strokes

So far, the CT afferent pathway has been the only afferent system to have a unique relationship with both stroking velocity and perceived pleasantness (Ackerley, 2022; Crucianelli et al., 2022), making it the prime candidate for mediating tactile processing of pleasant touch. Our findings showed that vibrotactile strokes, which target A*β* afferents, can induce a comparable sense of pleasantness to brush touch and confirm the results from Huisman et al. (2016). We further highlight differences with brush strokes by showing that both types of stroke were rated equally pleasant at lower velocities (0.5 and 1 cm/s), whereas brush strokes became more pleasant as velocity increased. The similarity of the U-shaped curves and of the pleasantness ratings at slow speed suggests that common mechanisms are at play, potentially related to tactile speed inference, even though the two types of stimulation elicit different neural responses.

The observed difference in pleasantness at high velocities for these stimulations could stem from multiple factors. For instance, the coefficient of friction and the surface roughness strongly modulate its pleasantness during tactile interaction (Gwosdow et al., 1986; Klöcker et al., 2013). Henceforth, it is likely that the intrinsic physical properties of the brush generated a different friction or roughness at high speeds, an effect that would not happen in the case of a motion illusion, in which only activation intervals change. Furthermore, we applied vibrotactile stimulation at a frequency of 120 Hz, which may interact with the perceived speed by eliciting a subjective sensation of faster tactile speed compared to a lower frequency. In view of the intensity curves in Experiment 1 (Figure 4C), it appears that the differences in perceived pleasantness between the two stimulations at low and high velocities cannot be explained by a common intensity-related coding mechanism.

### Impact of context and visuo-haptic speed integration

#### Invariance to companion velocity

The result that the pleasantness of a 3 cm/s stroke remains unaffected by its companion velocity is somewhat surprising, given that pleasantness has been shown to depend on the context (Croy et al., 2021; Löken et al., 2011; Triscoli et al., 2014). In our study, we expected the pleasant or unpleasant companion stimulation within an experimental block to be associated with a positive or negative emotional valence, which would bias the rating of the pleasantness of a 3 cm/s stroke (Convertino et al., 2024). Contrary to our hypothesis, it seems that participants judged the stimuli independently of each other, regardless of whether the stroke was performed using a brush or vibrations. This is somewhat surprising given that apparent motion does not activate CT afferents. There might have been habituation to the successive tactile stimuli that could have blunted the impact of the emotional valence, but no effect of block’s presentation order was found. The robustness of brush strokes at 3 cm/s is in line with the stable preference for that velocity, (Luong et al., 2017), which seems to also extend to vibrotactile strokes. Thus, the perceived tactile speed seems dominant over the psychological impact of the more pleasant or unpleasant context.

#### Impact of visuo-haptic discrepancy on pleasantness

Visual velocity did not affect the perceived pleasantness of soft brush, unlike vibrotactile strokes at 1 cm/s, which were more pleasant when presented in combination with faster, hence more pleasant visual velocities (Lee et al., 2018; Morrison et al., 2011; Walker et al., 2017; Willemse et al., 2016). A preliminary study suggested that the pleasantness of brush strokes can be affected by simultaneous visual strokes in virtual reality (Haraguchi & Kitazaki, 2022). In that study, 1 cm/s brush strokes were rated as less pleasant when visual velocities were slower or faster (Haraguchi & Kitazaki, 2022). However, that experiment mixed velocities differing by up to two orders of magnitude, and it did not preserve temporal synchrony at the start and end. Therefore, visuo-haptic discrepancy was very easy to notice and pleasantness ratings probably stemmed from the perception of congruence.

In our study, we limited visual speeds to deviations of up to 50% from the tactile reference, and we preserved the visuo-haptic synchrony of the stroke’s start and end. Thus, in most cases, participants were in most cases unable to perceive whether the visual and haptic stimulations were incongruent, which means that variations in stroke length that were induced by incongruent visual velocities were not detected. Notably, we found that the pleasantness slope induced by discrepant visual speeds mimicked the slope induced by actual variations of vibrotactile speed (Figure 7). This finding suggests that the pleasantness of vibrotactile apparent motion stems from velocity inference that can easily be manipulated. This inference would be derived from the underlying mechanisms that encode vibrotactile apparent motions. Future studies would need to test the impact of visuo-haptic integration in scenarios involving more significant discrepancies, which will enable to assess the additional influence of perceived discrepancy. This is particularly relevant in the case of a tactile stroke at 10 cm/s, in which the sensation of incongruence is stronger. Out of the ten visuo-tactile speeds presented, participants informally reported in a post-experiment interview that they have perceived four or five different speeds. This suggests that vision impacts somehow their conscious experience, but with limited effect on the report of pleasantness. In our study, a visuo-haptic stroke at 10 cm/s could have been more pleasant than one at 1 cm/s due to higher subjective congruence, but no difference in the ratings was observed. This observed lack of sensitivity to incongruity may stem from multisensory integration of pleasant touch being a largely unconscious process (Cleeremans & Sarrazin, 2007; Tononi et al., 2016). Indeed, the Integrated Information Theory suggests that complex cognitive tasks, such as assessing congruence in a scene, could be handled by feed-forward neuronal circuits that do not require the involvement of large integrative networks, such as the parietal cortex (Tononi et al., 2016).

Overall, our study shows that the pleasantness of both types of stimulation depends primarily on real or illusory tactile velocity. Visual input showed an impact on the perceived pleasantness of vibrotactile stimuli but not on physical brush strokes. This effect may be attributed either to the specific role of CT-activation for mediating pleasantness or to a more uncertain perception of vibrotactile velocity, making it more easily impacted by discrepant visual velocities. The respective contribution of visual inference and tactile stimulation strokes is still challenging and deserves additional investigations with a wider range of discrepant stimuli.

### Conclusion and perspectives

The velocity-dependent differences in pleasantness between vibrotactile and brush strokes indicate that the inverted-U shaped curve is not exclusively driven by CT afferents. This study also shows that the pleasantness of vibrotactile strokes is sensitive to visual velocity compared to brush strokes, but a potential limitation to consider is the visual brush we used to perform the strokes in the virtual environment. However, previous work showed that virtual female avatars and virtual feathers elicit similar pleasantness ratings (Seinfeld et al., 2022), suggesting that the effects reported here are likely robust across different visual representations. A strength of this study is the direct comparison of the two types of strokes on the same skin area in the same experiment. Direct comparison is particularly advantageous because pleasantness varies across skin body sites (Ackerley et al., 2014; Löken et al., 2011), and can be impacted by small differences in experimental conditions or the specific mindset of the tested group. To further investigate the generality of these findings, it would be interesting to use frequencies exactly matching those produced by a brush on the skin at different velocities to test whether a more natural frequency content could make vibrotactile pleasantness less prone to illusion. Likewise, it could be interesting to test the interplay between lower and higher velocities and lower and higher frequencies. Finally, measuring psychophysiological markers (Walker et al., 2022) such as the heart rate (Triscoli et al., 2017; Triscoli et al., 2017) or skin conductance (Etzi et al., 2018), induced by A*β*-touch or A*β* + CT touch, would be an effective method of finding distinguishable characteristics between these two distinct types of pleasant stimulation. Overall, these findings demonstrate that touch pleasantness arises from the integration of tactile cues, which extends beyond CT afferent tuning. They also highlight the importance of multisensory factors in shaping the perception of pleasant touch.

## Appendix

## Apparent haptic motion’s calculation

The ratio between *SOA* and *DoS* can give either a sensation of individual discrete point or continuous motion. Some studies have attempted to find an optimal *SOA* as a function of *DoS* (Israr & Abnousi, 2011; Israr & Poupyrev, 2011a), producing a general equation (Tactile Brush Algorithm) (Israr & Poupyrev, 2011b):

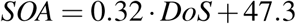

Considering the number of actuators and the fixed velocities (0.5, 1, 3, 10, and 30 cm/s) in our study, each value is computed as follows:

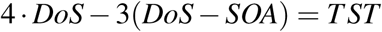

with *TST* defined as *v* = *d/TST*, the total stimulation time for each velocity. *SOA* value in both equations was resolved by selecting a *DoS* that is greater than *TST/*4 and less than *TST*. An iterative method was used to find a value *SOA* that approximates both equations, using various values *DoS*. Consequently, *SOA* was chosen as the value that minimizes the difference between the two calculated equations. These values are displayed in the table below.

Values of *DoS*, *SOA* and *TST* for each vibrotactile stroking stimulus.

| Velocity (cm/s) | 0.5 | 1 | 3 | 10 | 30 |
| --- | --- | --- | --- | --- | --- |
| **TST (s)** | 18 | 9 | 3 | 0.9 | 0.3 |
| **DoS (s)** | 9.11 | 4.52 | 1.45 | 0.39 | 0.08 |
| **SOA (s)** | 2.96 | 1.49 | 0.51 | 0.17 | 0.07 |

## Supplemental data

**S1)** Video showing congruent visuo-tactile brush strokes in first person perspective: https://doi.org/10.5281/zenodo.14632885.

## Author contributions statement

T-L S.L, G.B, M.A, and D.G conceived and designed research. D.V helps to implement the technical part. T-L S.L and D.V conducted the experiments, T-L S.L analysed data and T-L S.L, G.B, M.A and D.G interpreted the results of experiments. T-L S.L prepared figures and drafted manuscript, T-L S.L, G.B, M.A, and D.G edited and revised manuscript. T-L S.L, D.V, G.B, M.A, and D.G approved final version of manuscript.

## Additional information

### Competing interests

No conflicts of interest, financial or otherwise, are declared by the authors.

### Grants

This work was supported by a grant from the French National Research Agency (NeuroHCI) under Grant No. ANR-22-CE33-0006-01.

## Notes

https://doi.org/10.5281/zenodo.14632885

### Competing Interest Statement

The authors have declared no competing interest.

